## Supplementary Figures, Video Legends, and Tables for "Novel LOTUS-domain proteins are organizational hubs that recruit *C. elegans* Vasa to germ granules"

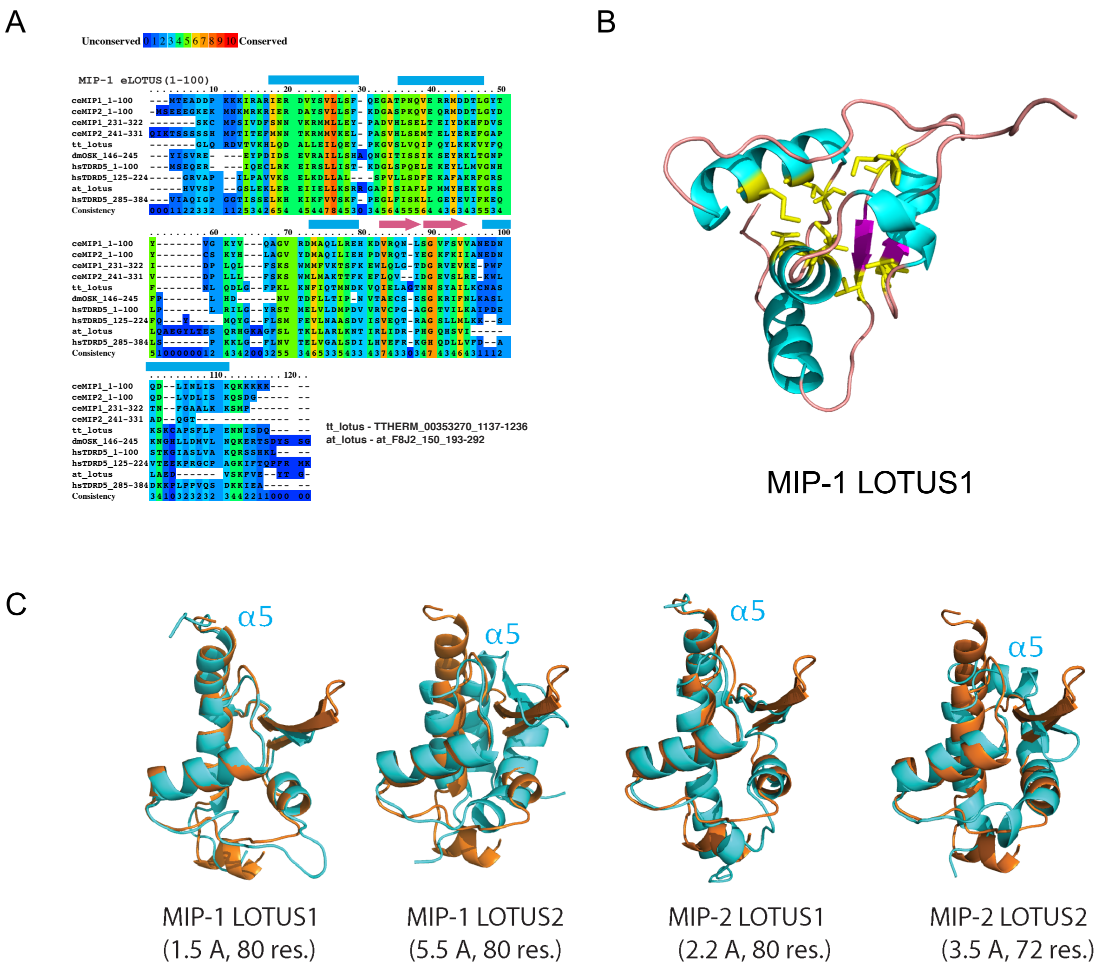


**Figure S1. Conserved residues in MIPs LOTUS domains are located in the central region of the structure. A,** Multiple sequence alignment showing conserved positions of predicted LOTUS domains in MIPs together with known LOTUS domains from other species: *T. thermophilus* B-box zinc finger protein with HTH OST-type domains (tt_lotus, residues 1137-1236); *D. melanogaster* Oskar (dmOSK, residues146-205); *H. sapiens* TDRD5 (hsTDRD5, residues 1-100, 125-224, and 285-384); and *A. thaliana* Zinc finger (CCCH-type) family protein with a HTH OST-type domain (at_lotus, residues 193- 292). **B**, Homology model of the MIP-1 LOTUS1 Three-dimensional structure. the most conserved residues (yellow, conservation score >5) are clustered in a conserved hydrophobic core. **C**, Structural overlaps of predicted MIP LOTUS domains (cyan) and the solved *Drosophila* Oskar eLOTUS domain (orange). RSMD values in Angstroms (Å) are also listed in **Table S2**.

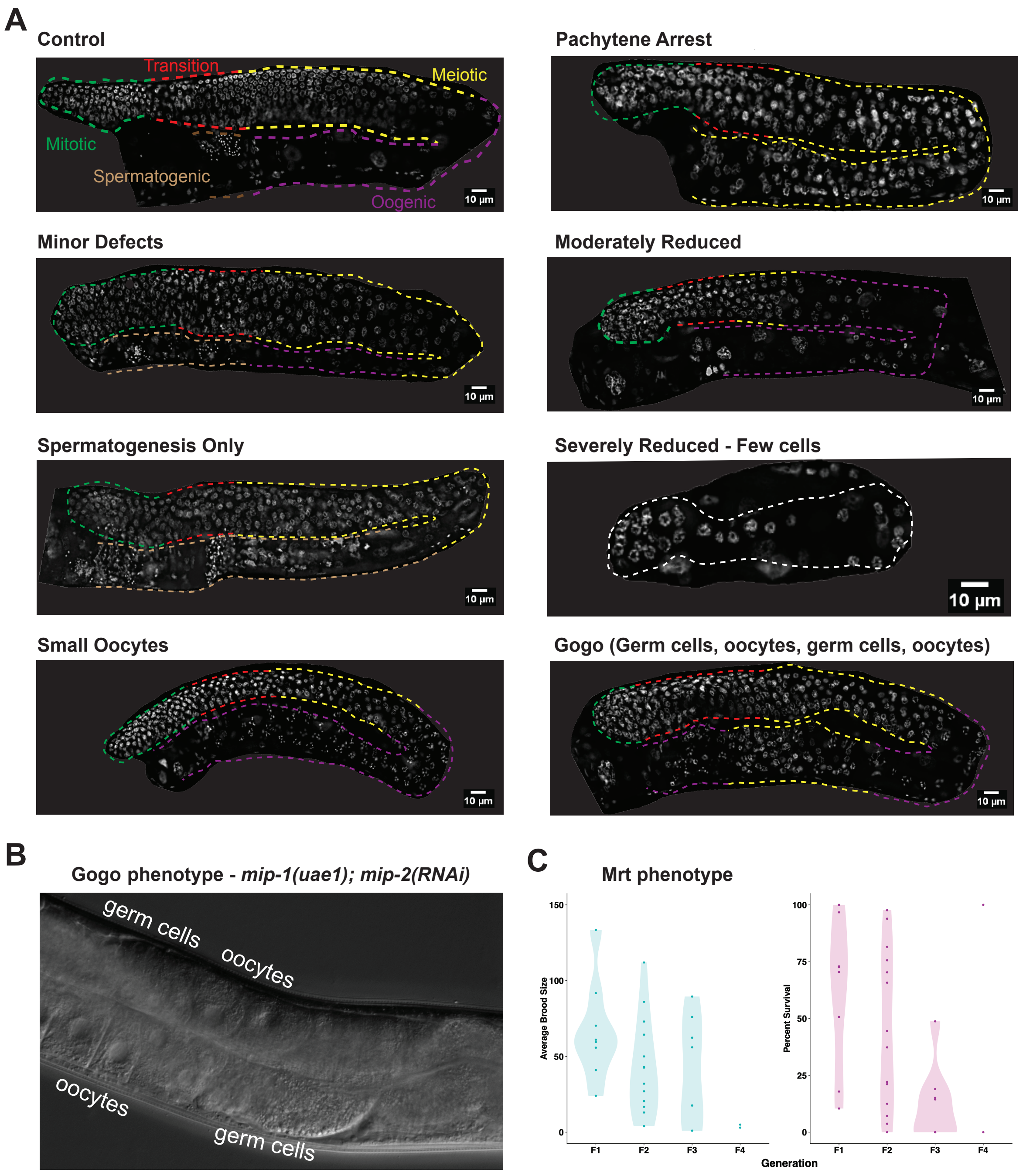


**Figure S2. Double *mip* null mutants show a similar array of germline phenotypes as double *mip* RNAi. A**, Phenotypes observed in the germ line of homozygous *mip-1(uae1);mip-2(uae-2)* double null mutants cultured at 25°C. Description of phenotypes in **Table S5**. **B,** DIC and images of a Gogo germ line. **C**, Quantification of average brood size and percentage survival of double *mip* null mutant progeny at 20°C across four successive generations. Lines show progressive loss of fertility, as evidenced by decreasing brood sizes per fertile mother, and higher embryonic lethality in the progeny, demonstrating a strong mortal germ line (Mrt) phenotype. Only two lines reach the F4 generation with at least one fertile parent, which produces an extremely low brood size. Data shown are the combined results of experiments that were started in the first generation and second generation after thawing (see Methods for details).

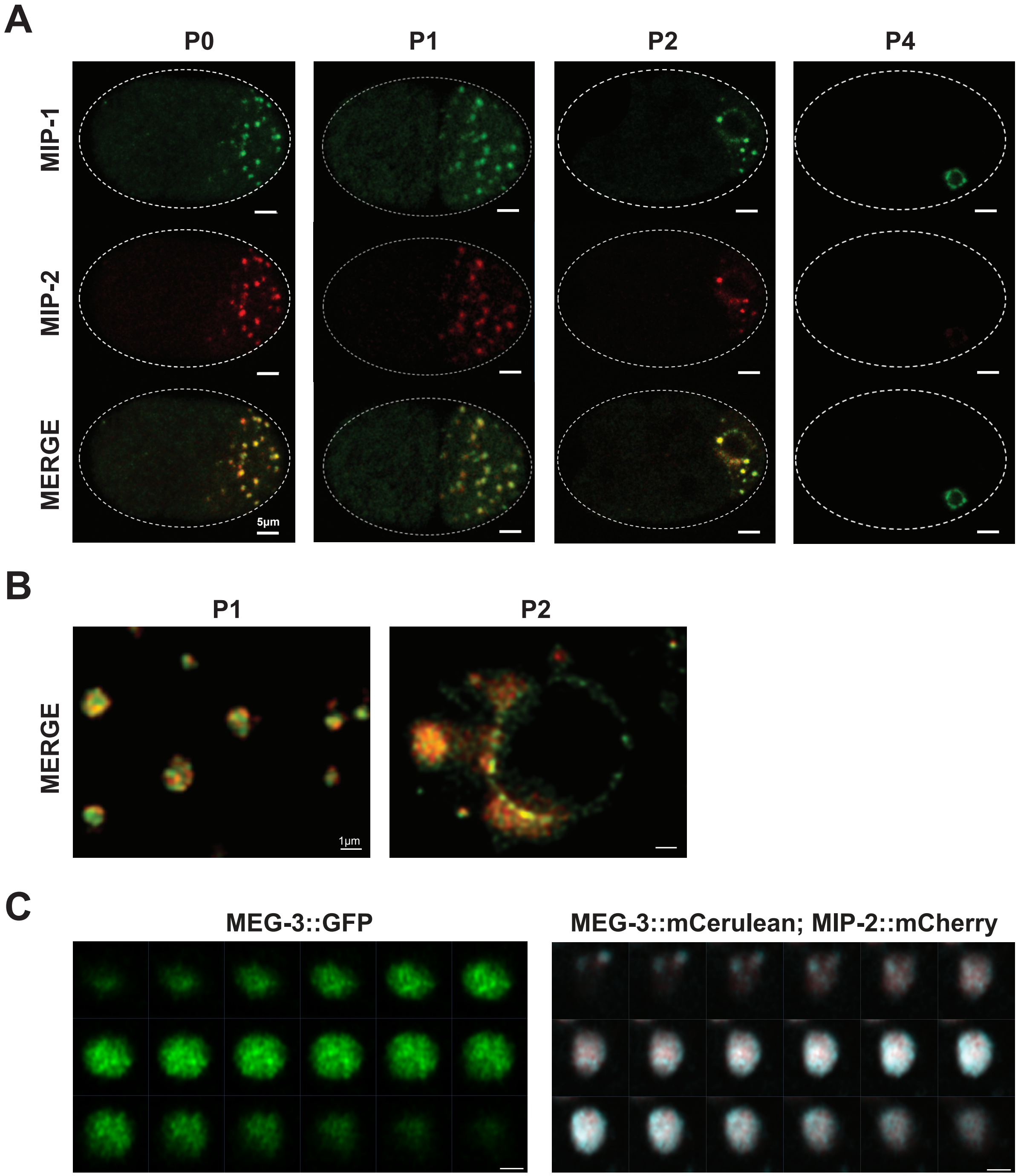


**Figure S3. MIPs and MEG-3 colocalize in live embryos.** Super-resolution micrographs of MIP-1 and MIP-2 granule segregation in live early embryos. **A,** MIP-1::GFP; MIP-2::mCherry embryos from 1-cell to 28-cells. **B**, Free-floating granules in the P1 cell of a two-cell embryo (left) and granule attachment to the nuclear membrane in the P2 cell of a 4-cell embryo (right)**.** **C,** Sequential images of Z-stacks through granules labeled with MEG-3::Cerulean and MIP-2::mCherry. Settings: 155nm Z-step, 2.8 µm Z-section. **D**, GFP::MEG-3. Settings: 75nm Z-step, 1.36µm Z-section. Scale bar: 1 µm.

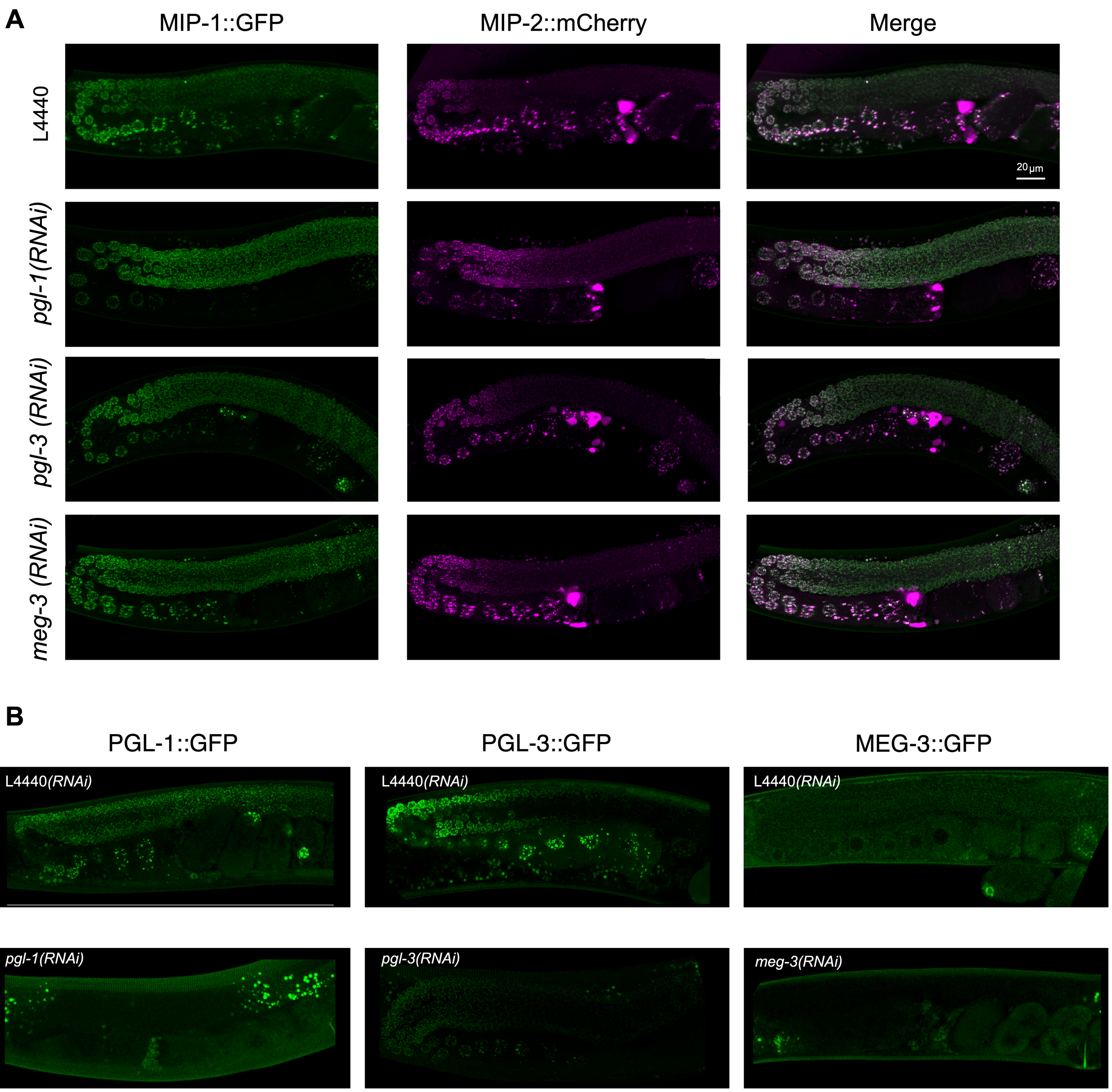


**Figure S4**. **Germline localization of MIP-1 and MIP-2 is not dependent on *pgl-1*, *pgl-3*, or *meg-3*.** Micrographs of MIP-1 and MIP-2 granules in germ lines from live animals. **A,** MIP-1::GFP;MIP-2::mCherry germ lines from animals treated with RNAi of *pgl-1*, *pgl-3*, *meg-3* and control RNAi (L4440). **B**, Control experiments to confirm effectiveness of the RNAi constructs used in the experiment shown in A. Scale bar: 20 μm.

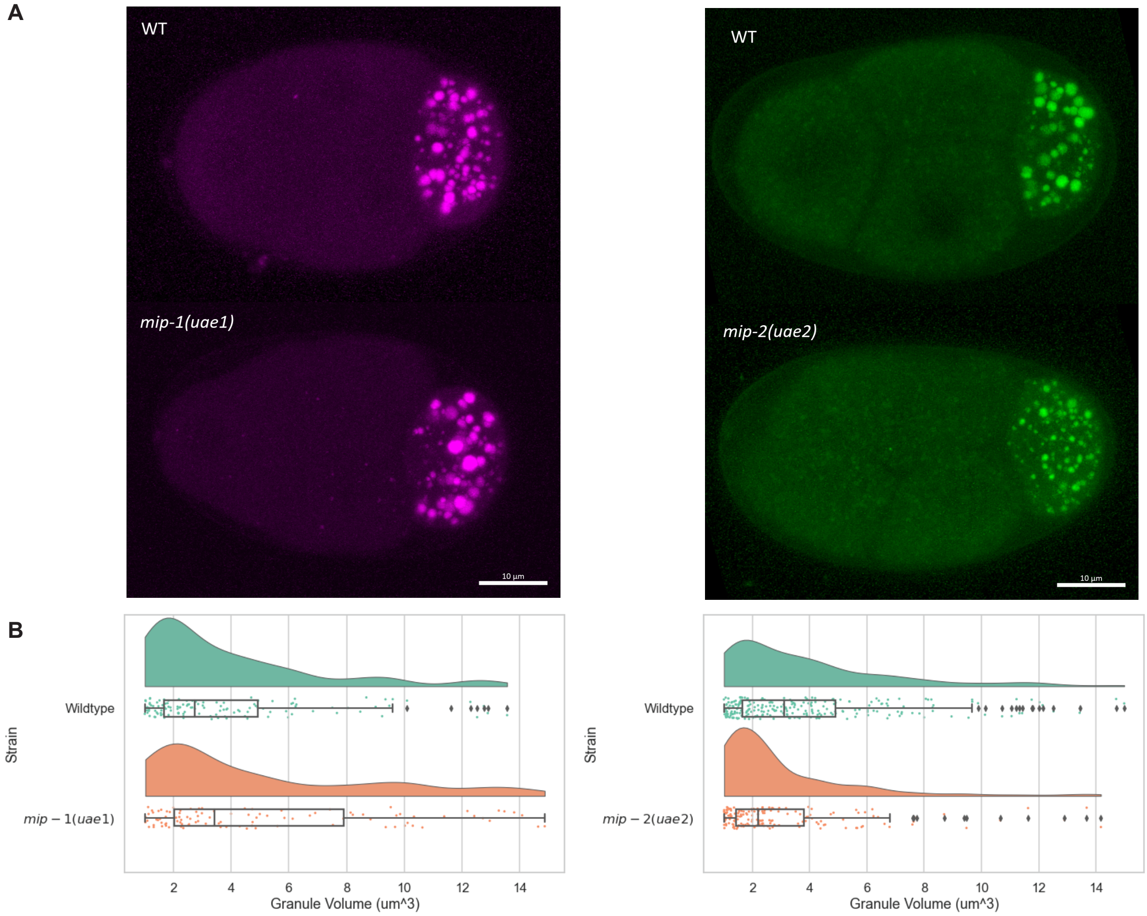


**Figure S5. MIPs affect each other's condensation in embryos. A,** MIP-2::mCherry four-cell embryos (left), and MIP-1::GFP four-cell embryos (right). **B,** Quantification of MIP granule volume in four-cell embryos comparing *mip* deletion strains with their corresponding controls. MIP-2 granules are larger in average when *mip-1* is deleted, and MIP-1 granules are smaller in average when *mip-2* is deleted.

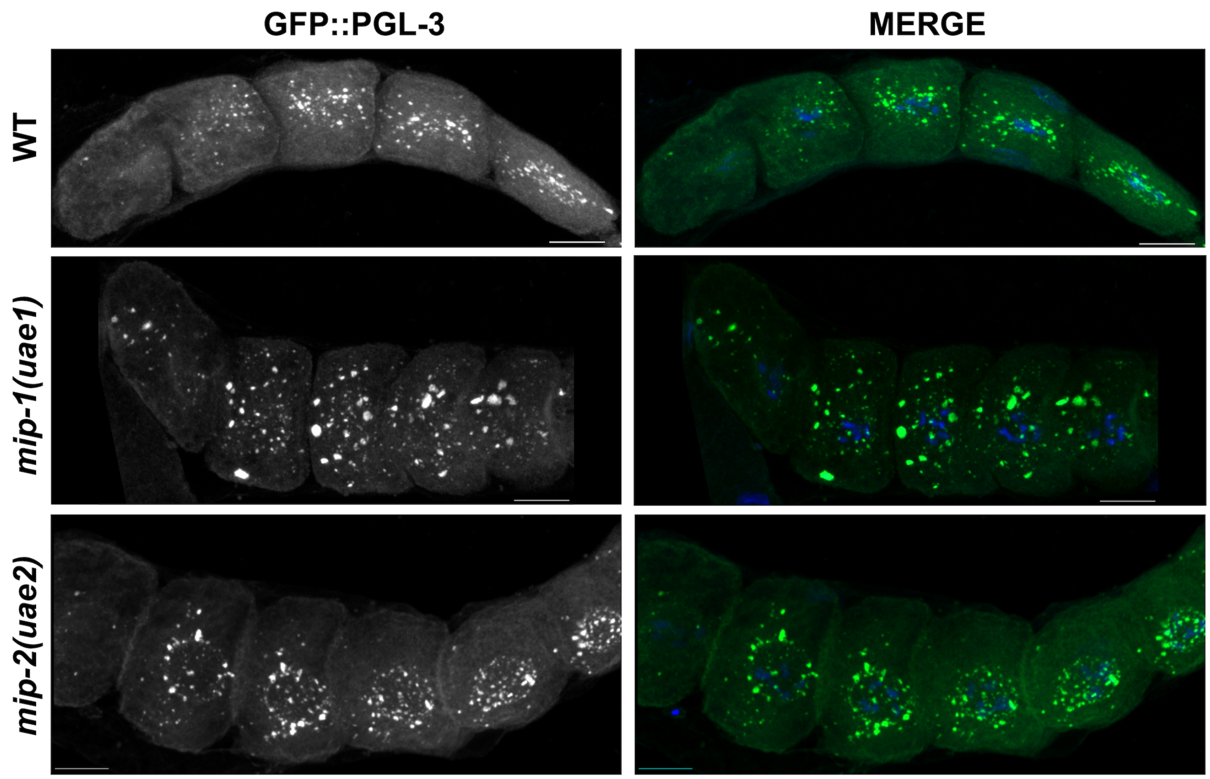


**Figure S6. MIP-1 and MIP-2 are required for the proper localization of PGL-3 granules.** Localization of GFP::PGL-3 in oocytes from adult dissected gonads of different genetic backgrounds: Wild-type strain (JH2017) (top), *mip-1(uae1)* null (strain GKC525) (middle), and *mip-2(uae2)* null (strain GKC548) (bottom). Panels labeled "MERGE" show overlay of GFP (green) and DAPI (DNA, blue) signals.

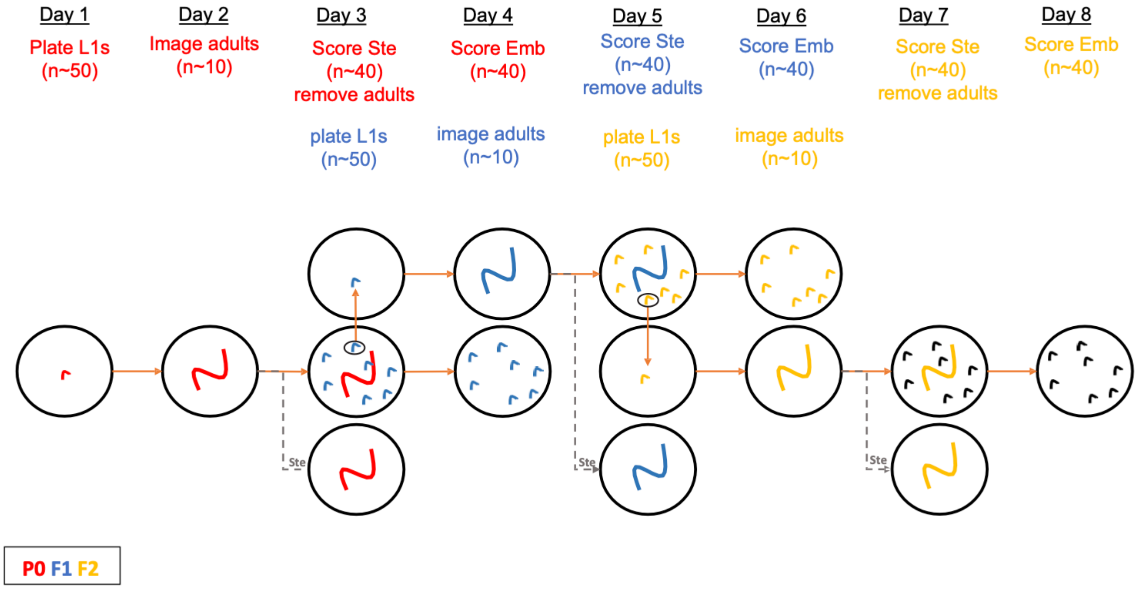


**Figure S7.** Protocol and scheduling for RNAi treatment and scoring of phenotypes for generation of data in Figure 2 (see also Materials and Methods).

#### Video Legends

**Video 1. Normal formation of PGL-3 granules is affected in the early embryo when *mips* are depleted.** Time lapse acquisition of the first two rounds of cell division in embryos from animals carrying GFP::PGL-3 and treated with L4440 control (top) or with *mip-1* and *mip-2* double RNAi (bottom).

**Video 2. Normal formation of GLH-1 granules is affected in the early embryo when *mips* are depleted.** Time lapse acquisition of the first two rounds of cell division in embryos from animals carrying GLH-1::GFP and treated with L4440 control (top) or with *mip-1* and *mip-2* double RNAi (bottom).

**Video 3. Normal formation of MEG-3 granules is affected in the early embryo when *mips* are depleted.** Time lapse acquisition of the first two rounds of cell division in embryos from animals carrying MEG-3::GFP and treated with L4440 control (top) or with *mip-1* and *mip-2* double RNAi (bottom).

**Video 4. Normal formation of MIPs granules is affected in the early embryo of a *meg-3* null mutant.** Time lapse acquisition of early embryonic development of embryos from animals carrying MIP-1::GFP and MIP-2::mCherry in a WT genetic background (top) and a *meg-3* null mutant background (bottom).

**Video 5. MEG-3 localizes to P granules in the embryo but not in the adult germ line.** Z-stack acquisition through a gonad arm including a few embryos in a strain carrying MEG-3::mCerulean and MIP-2::mCherry.

**Video 6. MIP-2 granule formation depends on *mip-1* in the early embryo.** Time-lapse acquisition of the first two rounds of cell division in embryos from animals carrying MIP-2::GFP in a WT genetic background (top) and a *mip-1* null mutant background (bottom).

**Video 7. MIP-1 granule formation depends on *mip-2* in the early embryo.** Time-lapse acquisition of the first two rounds of cell division in embryos from animals carrying MIP-1::GFP in a WT genetic background (top) and a *mip-2* null mutant background (bottom).

**Video 8. Title. Localization of GLH-1 granules is affected in germ cells when individual *mips* are removed.** Z-stack acquisition through a section of the gonad of animals carrying GLH-1::GFP in a WT background (left), a *mip-1* null background (middle), or a *mip-2* null background (right). Pachytene germ cells and oocytes are visible.

### Supplementary Tables

**Table S1. Proteins enriched in MEG-3::GFP pulldowns.** Proteins are sorted according to their *p*-value from the *t* SAM statistic as previously described (Chen et al., 2016). Data will become publicly available upon publication.

**Table S2. Similarity measures between predicted MIP LOTUS domain structures and LOTUS domains from other metazoans.** Protein domains: M1, MIP-1; M2, MIP-2; L1, LOTUS1; L2, LOTUS2. SeqID, sequence identity; GDT, Global Distance Test parameters for best templates; PDB ID, protein databank identifier of the best template; RMSD, backbone root mean square distances in Å, with number of aligned residues in parentheses. Published structures used for comparison: *D. melanogaster* Oskar (PBD ID 5nt7), *H. sapiens* TDRD5 (PBD ID 3s93).

| Protein domain | Coordinates | SeqID | GDT | PDB ID | RMSD (length) 5nt7 | 3s93 |
|---|---|---|---|---|---|---|
| M1 L1 | 1-101 | 16 | 33 | 5nt7 | 0.9 (80) | 3.7 (72) |
| M1 L2 | 227-314 | 10 | 50 | 2lh9 | 4.2 (72) | 1.6 (72) |
| M2 L1 | 1-104 | 14 | 51 | 5nt7 | 2.0 (80) | 1.4 (72) |
| M2 L2 | 243-327 | 16 | 57 | 3s93 | 3.3 (72) | 3.1 (80) |

**Table S3. Pairwise MIP LOTUS structural similarity analysis.** Values shown are backbone rmsd values in Å and number of aligned residues (in parentheses). M1, MIP-1, M2, MIP-2; L1, LOTUS1; L2, LOTUS2.

| Protein domain | M1 L2 | M2 L1 | M2 L2 |
|---|---|---|---|
| M1 L1 | 4.1 (72) | 3.0 (88) | 3.9 (80) |
| M1 L2 | | 1.7 (72) | 3.1 (80) |
| M2 L1 | | | 1.6 (80) |

**Table S4. Pairwise predicted binding affinities between MIP LOTUS domains and between MIP LOTUS domains and GLH-1**. Affinities are in kcal/mol. M1, MIP-1; M2, MIP-2; L1, LOTUS1; L2, LOTUS2. Predictions for GLH-1 binding considered only the helicase CTD domain. Predictions for combinations with no values given were highly unfavorable (>0 kcal/mol). Predicted binding affinities for the native *Drosophila* complexes: Oskar LOTUS homodimer = -54.5 kcal/mol; Oskar LOTUS—Vasa helicase complex = -53.5 kcal/mol.

| Protein/ domain | M1 L1 | M1 L2 | M2 L1 | M2 L2 |
|---|---|---|---|---|
| M1 L1 | | -16.7 | -21.1 | |
| M1 L2 | | | -5.3 | |
| M2 L1 | | | | |
| M2 L2 | | | | -6.1 |
| GLH-1 | -27.1 | | -4.9 | -0.8 |

**Table S5. Description of MIP depletion phenotypes.**

| Phenotype | Description |
|---|---|
| **Minor Defects** | No major or significant difference from germlines treated with the RNAi control (L4440). |
| **Reduced Germline** | There are many variants of this phenotype, having in common the overall reduction of multiple regions of the germline. There are often fewer oocytes and smaller pachytene regions. These animals are not always sterile. |
| **Abnormal Oocytes** | The size of some oocytes is significantly smaller than observed in control animals. These smaller oocytes tend to accumulate next to the spermatheca. The chromosome morphology in the oocytes tends to be abnormal and these animals make abnormal embryos as well. |
| **Spermatogenesis Only** | The germline produces only male gametes. Through late hermaphrodite adulthood, the germline fails to switch from a spermatogenic program to an oogenic program. |
| **Pachytene Arrest** | The germline nuclei do not progress in the meiotic division beyond the pachytene stage. The germline turn is present, and all nuclei beyond the turn have chromosomes with pachytene morphology. Often, there is also early entry into pachytene with reduced mitotic and transition zones. |
| **Few Germ Cells** | Germ cell populations are far fewer than normal. The germ cell number ranges from 2 to 14. The DNA morphology of these cells is abnormal. |

**Table S6. Strains produced and used in this study.**

| Name | Description | Genotype | Reference |
|---|---|---|---|
| GKC1 | *mip-1Δ* | *mip-1(uae1) III* | This study |
| GKC2 | *mip-2Δ* | *mip-2(uae2) V* | This study |
| GKC3 | *mip-1Δ; mip-2Δ* | *mip-1(uae1) III; mip-2(uae2) V* | This study |
| GKC6 | *meg-3Δ*; *mip-1::GFP*; *mip-2::mCherry* | *mip-1(knu191[mip-1::GFP]) III; mip-2(knu412[mip-2::mCherry]) V; meg-3(uae4) X* | This study |
| GKC501 | *mip-1::3xFLAG* | *mip-1(usa2[mip-1::3xFLAG]) III* | This study |
| GKC502 | *mip-2::3xFLAG* | *mip-2(usa3[mip-2::3xFLAG]) V* | This study |
| GKC509 | *lag-2p::GFP*; *his-58::mCherry*; *GFP::PH(PLC1δ1)* | *qIs153[lag-2p::MYR::GFP + ttx-3p:DsRed]; ltIs37[(pAA64) pie-1p::mCherry::his-58 + unc-119(+)] IV; ltIs38[pAA1; pie-1p::GFP::PH(PLC1δ1) + unc-119(+)]* | This study |
| GKC511 | *mip-2::GFP* | *mip-2(usa4[mip-2::GFP]) V* | This study |
| GKC518 | *mip-1::GFP; mip-2::mCherry* | *mip-1(knu191[mip-1::GFP]) III; mip-2(knu412[mip-2::mCherry]) V* | This study |
| GKC525 | *mip-1Δ*; *GFP::pgl-3* | *mip-1(uae1) unc-119(ed3)? III; axIs1464[pie-1p::GFP::pgl-3 ORF::pgl-3 3'utr + unc-119(+)]* | This study |
| GKC548 | *mip-2Δ*; *glh-1::GFP* | *glh-1(sam24[glh-1::gfp::3×FLAG]) I; mip-2(uae2) V* | This study |
| GKC551 | *mip-1Δ*; *glh-1::GFP* | *glh-1(sam24[glh-1::gfp::3×FLAG]) I; mip-1(uae1) III* | This study |
| GKC553 | *mip-1Δ*; *mip-2::mCherry* | *mip-1(uae1) III; mip-2(knu412[mip-2::mCherry])V* | This study |
| GKC554 | *mip-1::GFP; mip-2Δ* | *mip-1(knu191[mip-1::GFP]) III; mip-2(uae2) V* | This study |
| GKC555 | *mip-1Δ*; *mip-2Δ*; *glh-1::GFP* | *glh-1(sam24[glh-1::gfp::3×FLAG]) I; mip-1(uae1) III; mip-2(uae2) V* | This study |
| GKC564 | *mip-1::GFP* | *mip-1[knu191 - pNU696 (C-terminal GFP, loxP::unc-119(+)::loxP)] unc-119 (ed3) III* | This study |
| GKC565 | *mip-2::mCherry* | *mip-2 (knu412[mip-2::mCherry])* | This study |
| GKC566 | *mCherry::pgl-1; mip-1::GFP* | *mip-1(knu191[mip-1::GFP]) unc-119(ed3) III; tjIs15[pie-1::mCherry::pgl-1+unc-119(+)]* | This study |
| GKC567 | *mip-2Δ*; *GFP::pgl-3* | *unc-119(ed3) III; mip-2(uae2) V; axIs1464[pie-1p::GFP::pgl-3 ORF::pgl-3 3'utr + unc-119(+)]* | This study |
| DUP64 | *glh-1::GFP* | *glh-1(sam24[glh-1::gfp::3×FLAG]) I* | Andraloj K et al. *PLoS Genet* (2017) |
| SA180 | *mCherry::pgl-1* | *unc-119(ed3) III; tjIs15[pie-1::mCherry::pgl-1+unc-119(+)]* | Sugimoto A. (personal communication) |

| | | | |
|---|---|---|---|
| OD95 | *his-58:mCherry; GFP::PH(PLC1δ1)* | *unc-119(ed3) III; ltIs37[(pAA64) pie-1p::mCherry::his-58 + unc-119(+)] IV; ltIs38[pAA1;pie-1p::GFP::PH(PLC1δ1) + unc-119(+)]* | McNally et al., *J Cell Biol* (2006) |
| JH3016 | *GFP::meg-3* | *unc-119(ed3) III; axIs2076 [meg-3p::GFP::meg-3::meg-3 3'UTR + unc-119(+)]* | Wang JT, et al. *eLife* (2014) |
| JH3503 | *meg-3*::GFP | *meg-3(ax3054[meg-3::meGFP]) X* | Smith et al. *eLife* (2016) |
| JK4475 | *lag-2p::MYR::GFP* | *qIs153[lag-2p::MYR::GFP + ttx-3p:DsRed]* | Byrd DT, et al. *PLoS One* (2014) |
| JH2017 | *GFP::pgl-3* | *unc-119(ed3) III; axIs1464[pie-1p::GFP::pgl-3 ORF::pgl-3 3'utr + unc-119(+)]* | Gallo et al. *Science* (2010) |

**Table S7. Guide RNA sequences, repair templates, and screening primers for CRISPR strains produced in this study.**

| Target gene | Target Location (strand +/-) | Strain Desc-ription | Guide Sequence | Repair Template Sequence | Screening Primers (Forward / Reverse) |
|---|---|---|---|---|---|
| *dpy-10* | Exon 5/ (+) | Co-CRISPR screening tool | GCTACCATAGGCACCACGAG | CACTTGAACTTCAATACGGCAAGATGAGAATGAC TGGAAACCGTACCGCATGCGGTGCCTATGGTAGC GGAGCTTCACATGGCTTCAGACCAACAGCCTAT | |
| *mip-1* | 50nt upstream of C-terminus (+) | MIP-1::3xFLAG | TATCAATGCTGCGTTGCCGT | ACCACAACAATCGACTACATCTTCAATTGATAAT GAGTGTCTAGAAGCTATCAATGCTGCGTTGCCGT CGGATAAGGATAGTTGGGACTACAAAGACGATGA CGACAAGGACTACAAAGACGATGACGACAAGGAC TACAAAGACGATGACGACAAGTGATCGAAATTTT ACGTGCTTTAAAATATCCTGTTTATGTGTT | TCACCACAACAATCGACTACA / GCCAATCTTGACATGGCAGA |
| *mip-2* | C-terminus (+) | MIP-2::3xFLAG | GCACTGCTTCAACTACGCCT | TGAACTCCGATTTCTCGCCGAACATCTCCCAGGC GGACTACAAAGACGATGACGACAAGGACTACAAA GACGATGACGACAAGGACTACAAAGACGATGACG ACAAGTAGTTGAAGCAGTGCTCCTGACACGTATT TTATAATAT | ATTCCACCGTATCCCCGTAC / GGGTGTGAAAATACGGCCTC |
| *mip-1* | Full gene deletion (+/-) | *mip-1* null allele | N-term: GACATTCACTTGGCAAATGA C-term: TGCCGTCGGATAAGGATAGT | gatcaaattgtaaaatgttctctcagaagtgaca ttcacttggcaaATGTGAtcgaaattttacgtgc tttaaaatatcctgtttatgtgtttttgtaaaat tattttttaattag | CGTGCATCAGACTGGGTCTG / TTCGTGAATGACTCGCATCC |
| *mip-2* | Full gene deletion (+/-) | *mip-2* null allele | N-term: tgaaaaATGTCTGAAGAAGA C-term: GCACTGCTTCAACTACGCCT | gaatatttaaagtcattcaactgattgttttact gtttccagcatttgcGtgaaaaATGTAGttgaag cagtgctcctgacacgtattttataatatttatg ttttttttg | GCCACGATTTTGACATTTTTAAG / CGAAAATAGCGAAAATGGTTC |
| *meg-3* | Full gene deletion (+/-) | *mip-1* null allele | N-term: ACCGCTTGGGTAAGGTTTTG C-term: ttggtacaaTCATTGATCTC | atttatttcaatatttcattaagttttgattttt gcaggtATGTGAttgtaccaatttatatctatta cttgtagactatattgtatg | CAATATTTCATTAAGTTTTG / CATCCAGACAACAATATAATAC |

#### Table S8. Plasmid DNA constructs for *in vitro* pulldown experiments.

| Gene | Segment | Vector / tag | Antibiotic Resistance | Restriction Site (F/R) | Forward / Reverse Primer |
|---|---|---|---|---|---|
| *mip-1* | Full | pET-28-SUMO / SUMO-His | Kan | SacI / NotI | TGACGTTGAGCTCATGACGGAAGCTGACGATCCC / TGACGTTGCGGCCGCCCAACTATCCTTATCCGACGGC |
| *mip-1* | Full | pGEX-6p-1 / GST | Amp | SalI / NotI | TGACGTTGTCGACTCATGACGGAAGCTGACGATCCC / TGACGTTGCGGCCGCCCAACTATCCTTATCCGACGGC |
| *mip-1* | N-terminus | pET-28-SUMO / SUMO -His | Kan | SacI / NotI | TGACGTTGAGCTCATGACGGAAGCTGACGATCCC / TGACGTTGCGGCCGCCGGACGAACAACTGGTATTTG |
| *mip-1* | C-terminus | pET-28-SUMO / Sumo-His | Kan | SacI / NotI | TGACGTTGAGCTCATAGTGTCTGTCGCAGAAAGTATGAAAGG / TGACGTTGCGGCCGCCCAACTATCCTTATCCGACGGC |
| *mip-1* | N-terminus | pGEX-6p-1 / GST | Amp | SalI / NotI | GTCGACTCGAGCGGCCGCATATGACGGAAGCTGACGATCC / TGACGTTGCGGCCGCCGGACGAACAACTGGTATTTG |
| *mip-1* | C-terminus | pGEX-6p-1 / GST | Amp | NotI / NotI | TGACGTTGCGGCCGCATATAGTGTCTGTCGCAGAAAGTATGAAAGG / TGACGTTGCGGCCGCCCAACTATCCTTATCCGACGGC |
| *mip-1* | LOTUS 1 | pGEX-6p-1 / GST | Amp | SalI / NotI | TGACGTTGTCGACTCATGACGGAAGCTGACGATCCC / TGACGTTGCGGCCGCTTTCTTTTTTTTCTTCTGCTTTGAGATC |
| *mip-1* | LOTUS 2 | pGEX-6p-1 / GST | Amp | SalI / NotI | TGACGTTGTCGACTCTTGCAACCCGGGATTGACTCG / TGACGTTGCGGCCGCAGGAGCAGGTGGCATTGATTTC |
| *mip-2* | Full-length | pET-28-SUMO / SUMO -His | Kan | SacI / NotI | TGACGTTGAGCTC atgtctgaagaagaaggcaaagaaaaaatg / TGACGTTGCGGCCGCCGCCTGGGAGATGTTCGGCG |
| *mip-2* | Full-length | pGEX-6p-1 / GST | Amp | XmaI (2x) / NotI | TGACGTTCCCGGGCCCGGGTatgtctgaagaagaaggcaaagaaaaaatg / TGACGTTGCGGCCGCCGCCTGGGAGATGTTCGGCG |
| *mip-2* | N-terminus | pET-28-SUMO / SUMO -His | Kan | BamHI / SalI | TGACGTTGGATCCatgtctgaagaagaaggcaaagaaaaaatg / TGACGTTGTCGACGTATCTAAACCCTTGTGATATTGGTAGCTG |
| *mip-2* | C-terminus | pET-28-SUMO / SUMO -His | Kan | SacI/ NotI | TGACGTTGAGCTCcaggatcctaactcaaagccgaaag / TGACGTTGCGGCCGCCGCCTGGGAGATGTTCGGCG |
| *mip-2* | C-terminus | pGEX-6p-1 / GST | Amp | NotI/ NotI | TGACGTTGCGGCCGCATCAGGATCCTAACTCAAAGCCGAAAG / TGACGTTGCGGCCGCCGCCTGGGAGATGTTCGGCG |
| *mip-2* | LOTUS 1 | pGEX-6p-1 / GST | Amp | SalI/ NotI | TGACGTTGTCGACTCATGTCTGAAGAAGAAGGCAAAGAAAAAATG / TGACGTTGCGGCCGCTCCGTCGCTCTGCTTCGAGATC |
| *mip-2* | LOTUS 2 | pGEX-6p-1 / GST | Amp | SalI/ NotI | TGACGTTGTCGACTCCCTGTCAAGAAGCATGTGC / TGACGTTGCGGCCGCAGCAACAGTTGTAGCTTCAAC |
| *glh-1* | Full-length | pGEX-6p-2 / GST | Amp | SalI/ NotI | TGACGTTGTCGACTCATGTCTGATGGTTGGAGTGATAGCG / TGACGTTGCGGCCGCCCAGCCTTCTTCATCTTGAGGG |
| *glh-1* | Helicase domains | pGEX-6p-2 / GST | Amp | SalI/ NotI | TGACGTTGTCGACTCGGTGTTGAAGGAGAAGGACCTAAG / TGACGTTGCGGCCGCCCAGCCTTCTTCATCTTGAGGG |
